# Dark Control: Towards a Unified Account of Default Mode Function by Markov Decision Processes

**DOI:** 10.1101/148890

**Authors:** Elvis Dohmatob, Guillaume Dumas, Danilo Bzdok

**Affiliations:** INRIA, Parietal Team, Saclay, France; Neurospin, CEA, Gif-sur-Yvette, France; Department of Psychiatry, Psychotherapy and Psychosomatics, RWTH Aachen University, Aachen, Germany; JARA-BRAIN, Jülich-Aachen Research Alliance, Germany; Institut Pasteur, Human Genetics and Cognitive Functions Unit, Paris, France; CNRS UMR 3571 Genes, Synapses and Cognition, Institut Pasteur, Paris, France; University Paris Diderot, Sorbonne Paris Cité, Paris, France; Centre de Bioinformatique, Biostatistique et Biologie Intégrative, Paris, France

**Keywords:** systems neuroscience, artificial intelligence, reinforcement learning, mind-wandering

## Abstract

The default mode network (DMN) is believed to subserve the baseline mental activity in humans. Its highest energy consumption compared to other brain networks and its intimate coupling with conscious awareness are both pointing to an overarching function. Many research streams speak in favor of an evolutionarily adaptive role in envisioning experience to anticipate the future. In the present work, we propose a *process model* that tries to explain *how* the DMN may implement continuous evaluation and prediction of the environment to guide behavior. Specifically, we answer the question whether the neurobiological properties of the DMN collectively provide the computational building blocks necessary for a Markov Decision Process. We argue that our formal account of DMN function naturally accommodates as special cases previous interpretations based on (1) predictive coding, (2) semantic associations, and (3) a sentinel role. Moreover, this process model for the neural optimization of complex behavior in the DMN offers parsimonious explanations for recent experimental findings in animals and humans.

## 1 Introduction

In the absence of external stimulation, the human brain is not at rest. At the turn to the 21st century, brain-imaging may have been the first technique to allow for the discovery of a unique brain network that would subserve baseline mental activities (Raichle et al., 2001; Buckner et al., 2008; Bzdok and Eickhoff, 2015). The “default mode network” (DMN) continues to metabolize large quantities of oxygen and glucose energy to maintain neuronal computation during free-ranging thought (Kenet et al., 2003; Fiser et al., 2004). The baseline energy demand is only weakly modulated at the onset of defined psychological tasks (Gusnard and Raichle, 2001). At its opposite, during sleep, the decoupling of brain structures discarded the idea of the DMN being only a passive network resonance and rather supported an important role in sustaining conscious awareness (Horovitz et al., 2009).

This *dark matter of brain physiology* (Raichle, 2006) begs the question of the biological purpose underlying DMN activity. Despite observation of similar large-scale networks of co-varying spontaneous activity in electrophysiological investigations (De Pasquale et al., 2010; Brookes et al., 2011; Baker et al., 2014), the link between the fMRI BOLD signal and population-level neural activity is still unclear. If those frequency-specific electrophysiological correlations are proposed as complementary to those observed with BOLD (Hipp and Siegel, 2015), their role in DMN function remains elusive (Maldjian et al., 2014).

What has early been described as the “stream of consciousness” in psychology (James, 1890) found a potential neurobiological manifestation in the DMN (Shulman et al., 1997; Raichle et al., 2001). We propose that this set of some of the most advanced regions in the association cortex (Mesulam, 1998; Margulies et al., 2016b) are responsible for higher-order control of human behavior. Our functional account follows the notion of “a hierarchy of brain systems with the DMN at the top and the salience and dorsal attention systems at intermediate levels, above thalamic and unimodal sensory cortex” (Carhart-Harris and Friston, 2010).

### 1.1 Towards a formal account of default mode function: higher-order control of the organism

The network nodes that compose the human DMN are responsible for extended parts of the baseline neural activity, which typically decreases when engaged in controlled psychological experiments (Gusnard and Raichle, 2001). The standard mode of neural information maintenance and manipulation has been argued to mediate evolutionarily conserved functions (Brown, 1914; Binder et al., 1999; Buzsáki, 2006). Today, many psychologists and neuroscientists believe that the DMN implements some form of probabilistic estimation of past, hypothetical, and future events (Fox et al., 2005; Hassabis et al., 2007; Schacter et al., 2007; Binder et al., 2009; Buckner et al., 2008; Spreng et al., 2009). This brain network might have emerged to continuously predict the environment using mental imagery as an evolutionary advantage (Suddendorf and Corballis, 2007). However, information processing in the DMN has also repeatedly been shown to directly impact human behavior. Goal-directed task performance improved with decreased activity in default mode regions (Weissman et al., 2006) and increased DMN activity was linked to more task-independent, yet sometimes useful thoughts (Mason et al., 2007; Seli et al., 2016). Gaining insight into DMN function is particularly challenging because this brain network appears to simultaneously modulate perception-action cycles in the present and to support mental travel across time, space, and content domains (Boyer, 2008).

The present work adopts the perspective of a human *agent* faced with the choice of the next actions and guided by outcomes of really happened, hypothetically imagined, and expected futures to optimize behavioral performance. Formally, a particularly attractive framework to describe, quantify, and predict intelligent systems, such as the brain, is proposed to be the combination of control theory and reinforcement learning (RL). An intelligent agent improves the interaction with the environment by continuously updating its computation of value estimates and action predispositions through integration of feedback outcomes. That is, “[agents], with their actions, modify the environment and in doing so partially determine their next stimuli, in particular stimuli that are necessary for triggering the next action” (Pezzulo, 2011). Agents with other behavioral policies therefore sample different distributions of action-perception trajectories (Ghavamzadeh et al., 2015). Henceforth, *control* refers to the influence that an agent exerts when interacting with the environment to reach preferred states.

Psychologically, the more the ongoing executed task is unknown and unpracticed, the less stimulus-independent thoughts occur (Filler and Giambra, 1973; Teasdale et al., 1995; Christoff et al., 2016). Conversely, it has been empirically shown that, the more the world is easy to foresee, the more human mental activity becomes detached from the actual sensory environment (Antrobus et al., 1966; Pope and Singer, 1978; Mason et al., 2007; Weissman et al., 2006). Without requiring explicit awareness, these “offline” processes may contribute to optimizing control of the organism. We formalize a *policy matrix* to capture the space of possible actions that the agent can perform on the environment given the current state. A *value function* maps environmental objects and events (i.e., states) to expected rewards. Switching between states reduces to a sequential processing model. Informed by outcomes of performed actions, neural computation reflected in DMN dynamics could be constantly shaped by prediction error through feedback loops. The present computational account of DMN function will be described in the mathematical framework of Markov Decision Processes (MDP). MDPs specifically formalize decision making in stochastic contexts with reward feedback.

Such an RL account of DMN function can naturally embed human behavior into the tension between exploitative action with immediate gains and exploratory action with longer-term gratification. We argue that DMN implication in many of the most advanced human capacities can be recast as prediction error minimization informed by internally generated probabilistic simulations - “covert forms of action and perception” (Pezzulo, 2011) -, allowing maximization of action outcomes across multiple time-scales. Such a purposeful optimization objective may be solved by a stochastic approximation based on a brain implementation of Markov Chain Monte Carlo (MCMC) sampling. Even (unavoidably) imperfect memory recall, Even necessarily imperfect memory recall, random day-time mind-wandering, and seemingly arbitrary dreams during sleep may provide randomly sampled blocks of pseudo-experience instrumental to iteratively optimize the behavior of the organism.

Evidence from computational modeling of human behavior (Körding and Wolpert, 2004) and cell recording experiments in ferrets (Fiser et al., 2004) suggest that the brain is largely dedicated to “the development and maintenance of [a] probabilistic model of anticipated events” (Raichle and Gusnard, 2005). The present paper proposes a process model that satisfies this previously proposed contention. We also contribute to the discussion of DMN function by providing some of the first empirical evidence that morphological variability in DMN regions is linked to the reward circuitry (Fig. 2), thus linking two literatures with currently scarce cross-references. Finally, we detail how our process model relates to previous accounts of DMN function and we derive explicit hypotheses to be tested in future neuroscience experiments. At this stage, we emphasize the importance of differentiating which levels of observation are at play in the present account. A process model is not solely intended to capture behavior of the agent, such as cognitive accounts of DMN function, but also the neurocomputational specifics of the agent. Henceforth, we will use “inference” when describing aspects of the statistical model, “prediction” when referring to the neurobiological implementation, and words like “forecast” or “forsee” when referring to the behavior of the agent.

## 2 Known neurobiological properties of the default mode network

We begin by a neurobiological deconstruction of the DMN based on experimental findings in the neuroscience literature. This walkthrough across main regions of the DMN will outline the individual functional profiles, paving the way for their algorithmic interpretation in our formal account (section 3).

### 2.1 The posteromedial cortex: global monitoring and information integration

The midline structures of the human DMN, including the posteromedial cortex (PMC) and the medial prefrontal cortex (mPFC), are probably responsible for highest turn-overs of energy consumption (Raichle et al., 2001; Gusnard and Raichle, 2001). These metabolic characteristics go hand-in-hand with brain-imaging findings that suggested the PMC and mPFC to potentially represent the functional core of the DMN (Andrews-Hanna et al., 2010; Hagmann et al., 2008).

Normal and disturbed metabolic fluctuations in the human PMC have been closely related to changes of conscious awareness (Cavanna and Trimble, 2006). Indeed, the PMC matures relatively late (i.e., myelination) during postnatal development in monkeys (Goldman-Rakic, 1987), which is generally considered to be a sign of evolutionary sophistication. This DMN region has long been speculated to reflect constant computation of environmental statistics and its internal representation as an inner “mind’s eye” (Cavanna and Trimble, 2006; Leech and Sharp, 2014). For instance, Bálint’s syndrome is a neurological disorder of conscious awareness that results from medial damage in the parietal cortex (Bálint et al., 1909). Such neurological patients are plagued by an inability to bind various individual features of the visual environment into an integrated whole (i.e., simultanagnosia) as well as an inability to direct action towards currently unattended environmental objects (i.e., optic ataxia). This dysfunction can be viewed as a high-level impairment in gathering information about alternative objects (i.e., exploration) as well as leveraging these environmental opportunities (i.e., exploitation). Congruently, the human PMC was coupled in two different functional connectivity analyses (Bzdok et al., 2015) with the amygdala, involved in significance evaluation, and the nucleus accumbens (NAc), involved in reward evaluation. Specifically, among all parts of the PMC, the ventral posterior cingulate cortex was most connected to the laterobasal nuclei group of the amygdala (Bzdok et al., 2015). This amygdalar subregion has been proposed to continuously scan environmental input for biological relevance assessment (Bzdok et al., 2013a; Ghods-Sharifi et al., 2009; Baxter and Murray, 2002).

The putative role of the PMC in continuous abstract integration of environmental relevance and ensuing top-level guidance of action on the environment is supported by many neuroscience experiments. Electrophysiological recordings in animals implicated PMC neurons in strategic decision making (Pearson et al., 2009), risk assessment (McCoy and Platt, 2005), outcome-dependent behavioral modulation (Hayden et al., 2009), as well as approach-avoidance behavior (Vann et al., 2009). Neuron spiking activity in the PMC allowed distinguishing whether a monkey would pursue an exploratory or exploitative behavioral strategy during food foraging (Pearson et al., 2009). Further, single-cell recordings in the monkey PMC demonstrated this brain region’s sensitivity to subjective target utility (McCoy and Platt, 2005) and integration across individual decision-making instances (Pearson et al., 2009). This DMN region encoded the preference for or aversion to options with uncertain reward outcomes and its neural spiking activity was more associated with subjectively perceived relevance of a chosen object than by its actual value, based on an “internal currency of value” (McCoy and Platt, 2005). In fact, direct stimulation of PMC neurons in monkeys promoted exploratory actions, which would otherwise be shunned (Hayden et al., 2008). Graded changes in firing rates of PMC neurons indicated changes in upcoming choice trials, while their neural patterns were distinct from neuronal spike firings that indicated choosing either option. Similarly in humans, the DMN has been shown to gather and integrate information over different parts of auditory narratives in an fMRI study (Simony et al., 2016).

Moreover, the retrosplenial portion of the PMC could support representation of action possibilities and evaluation of reward outcomes by integrating information from memory recall and different perspective frames. Regarding memory recall, retrosplenial damage has been consistently associated with anterograde and retrograde memory impairments of various kinds of sensory information in animals and humans (Vann et al., 2009). Regarding perspective frames, the retrosplenial subregion of the PMC has been proposed to mediate between the organism’s egocentric (i.e., focused on external sensory environment) and allocentric (i.e., focused on internal world knowledge) viewpoints in animals and humans (Epstein, 2008; Burgess, 2008; Valiquette and McNamara, 2007).

Consequently, the PMC may contribute to overall DMN function by monitoring the subjective outcomes of possible actions and integrating that information with memory and perspective frames into short and longer-term behavioral agendas. Estimated value, found to differs across individuals, might enrich statistical assessment of the environment to map and predict delayed reward opportunities in the future. In doing so, the PMC may continuously adapt the organism to changes in both the external environment and its internal representation to enable strategic behavior.

### 2.2 The prefrontal cortex: action consideration and stimulus-value association

Analogous to the PMC, the dorsomedial PFC (dmPFC) of the DMN is believed to subserve multi-sensory processes across time, space, and content domains to exert top-level control on behavior. Comparing to the PMC, however, dmPFC function may be closer to a “mental sketchpad” (Goldman-Rakic et al., 1996), as this DMN part potentially subserves the de-novo construction and manipulation of meaning representations instructed by stored semantics and memories (Bzdok et al., 2013c). The dmPFC may subserve representation and assessment of one’s own and other individuals’ action considerations. Generally, neurological patients with tissue damage in the prefrontal cortex are known to struggle with adaptation to new stimuli and events (Stuss and Benson, 1986). Specifically, neural activity in the human dmPFC reflected expectations about other peoples’ actions and outcomes of these predictions. Neural activity in the dmPFC indeed explained the performance decline of inferring other peoples’ thoughts in aging humans (Moran et al., 2012). Certain dmPFC neurons in macaque monkeys exhibited a preference for processing others’, rather than own, action with fine-grained adjustment of contextual aspects (Yoshida et al., 2010).

Comparing to the dmPFC, the vmPFC is probably more specifically devoted to subjective value evaluation and risk estimation of relevant environmental stimuli (Fig. 1 and 2). The ventromedial prefrontal DMN may subserve adaptive behavior by bottom-up-driven processing of what matters now, drawing on sophisticated value representations (Kringelbach and Rolls, 2004; O’Doherty et al., 2015). Quantitative lesion findings across 344 human individuals confirmed a substantial impairment in value-based action choice (Gläscher et al., 2012). Indeed, this DMN region is preferentially connected with reward-related and limbic regions. The vmPFC is well known to have direct connections with the NAc in axonal tracing studies in monkeys (Haber et al., 1995). Congruently, the gray-matter volume of the vmPFC and NAc correlated with indices of value-guided behavior and reward attitudes in humans (Lebreton et al., 2009). NAc activity is further thought to reflect reward prediction signals from dopaminergic neurotransmitter pathways (Schultz, 1998) that not only channel action towards basic survival needs but also enable more abstract reward processings, and thus perhaps RL, in humans (O’Doherty et al., 2015).

**Fig. 1.**
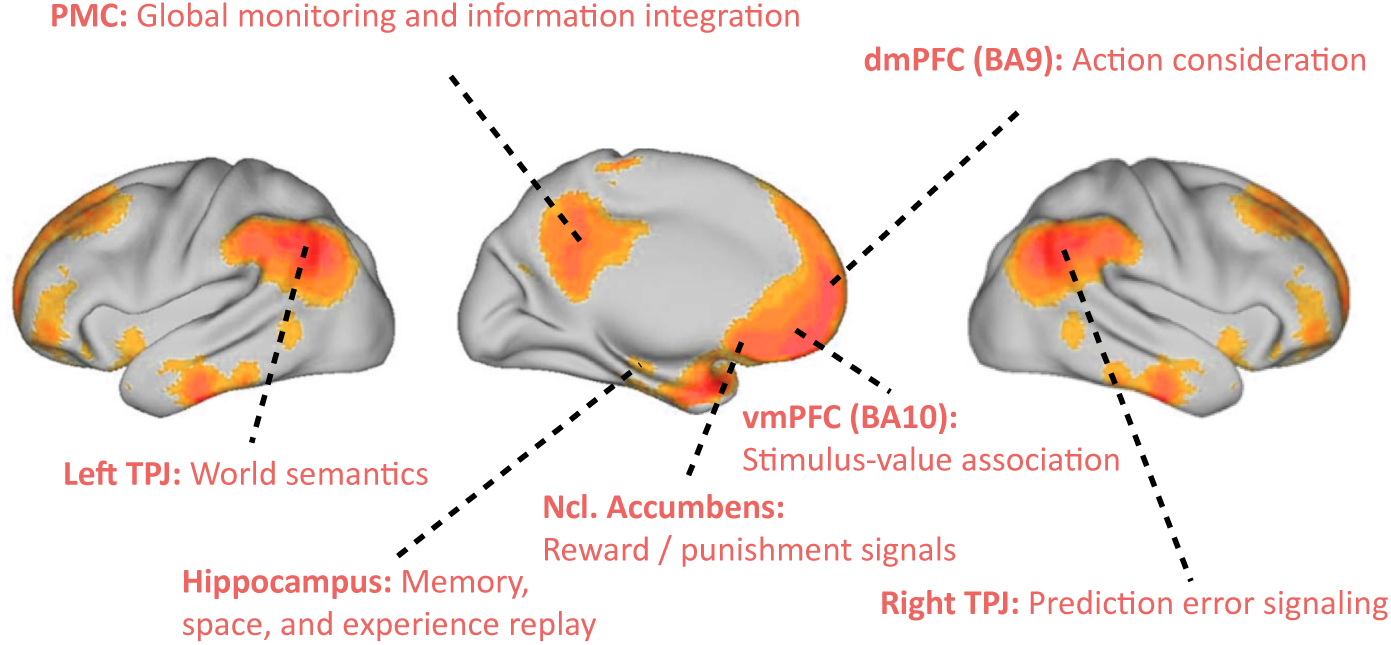
Default mode network: key functions. Neurobiological overview of the DMN with its major constituent parts and the associated functional roles relevant in our functional interpretation.

**Fig. 2.**
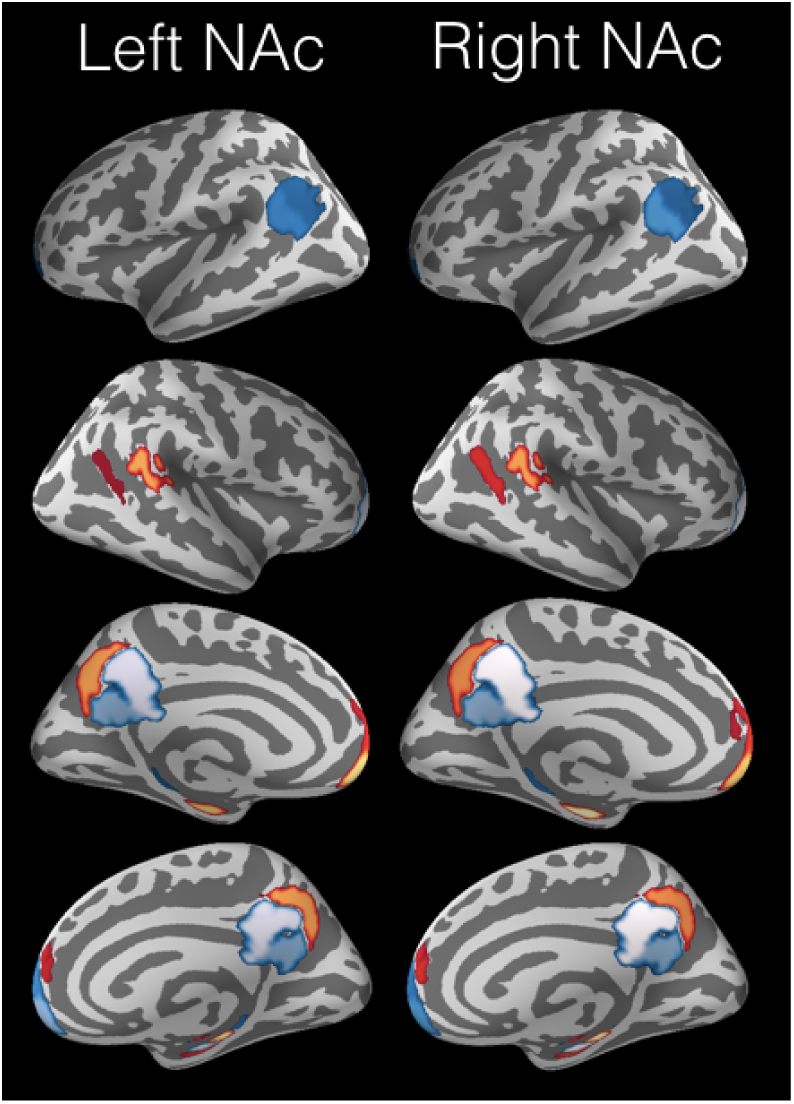
Morphological coupling between reward system and default mode network. Based on 9,932 human subjects from the UK Biobank, inter-individual differences in left NAc volume (*R*^*2*^ = 0.11±0.02) and right NAc volume (*R*^*2*^ = 0.14 ± 0.02) could be predicted from volume in the DMN regions. These out-of-sample generalization performances were obtained from support vector regression applied to normalized region volumes in the DMN in a 10-fold cross-validation procedure. Consistent for the left and right reward system, NAc volume in a given subject is positively coupled with the vmPFC and HC. The colors are indicative of the (red = positive, blue = negative) and relative importance (the lighter the higher) of the regression coefficients. The code for reproduction and visualization: www.github.com/banilo/darkcontrol_PCB2018.

Consistently, diffusion MRI tractography in humans and monkeys (Croxson et al., 2005) quantified the NAc to be more connected to the vmPFC than dmPFC in both species. Two different functional connectivity analyses in humans also revealed strong vmPFC connections with the NAc, hippocampus (HC), and PMC (Bzdok et al., 2015). In line with these connectivity findings in animals and humans, the vmPFC is often proposed to represent triggered emotional and motivational states (Damasio et al., 1996). Such real or imagined arousal states could be mapped in the vmPFC as a bioregulatory disposition influencing cognition and decision making. In neuroeconomic studies of human decision making, the vmPFC consistently reflects an individuals subjective value predictions (Behrens et al., 2008), which may also explain why performance within and across participants was reported to relate to state encoding in the vmPFC (Schuck et al., 2016). Such a “cognitive map” of the action space was argued to encode the current task state even when states are unobservable from the sensory environment.

### 2.3 The hippocampus: memory, space, and experience replay

The DMN midline has close functional links with the HC in the medial temporal lobe (Vincent et al., 2006; Shannon et al., 2013) a region long known to be involved in memory operations and spatial navigation in animals and humans. While the HC is traditionally believed to allow recalling past experience, there is now increasing evidence for an important role in constructing mental models in general (Zeidman and Maguire, 2016; Schacter et al., 2007; Gelbard-Sagiv et al., 2008; Javadi et al., 2017; Boyer, 2008). Its recursive anatomical architecture may be specifically designed to allow reconstructing entire sequences of experience from memory fragments. Indeed, hippocampal damage was not only associated with an impairment in re-experiencing the past (i.e., amnesia), but also forecasting of one’s own future and imagination of experiences more broadly (Hassabis et al., 2007).

Mental scenes created by neurological patients with HC lesion exposed a lack of spatial integrity, richness in detail, and overall coherence. Single-cell recordings in the animal HC revealed constantly active neuronal populations whose firing coincided with specific locations in space during environmental navigation. Indeed, when an animal is choosing between alternative paths, the corresponding neuronal populations in the HC spike one after another (Johnson and Redish, 2007). Such neuronal patterns in the HC appear to directly indicate upcoming behavior, such as in planning navigational trajectories (Pfeiffer and Foster, 2013) and memory consolidation of choice relevance (De Lavilléon et al., 2015). Congruently, London taxi drivers, humans with high performance in forecasting spatial navigation, were shown to exhibit increased gray-matter volume in the HC (Maguire et al., 2000).

There is hence increasing evidence that HC function extends beyond simple forms of encoding and reconstruction of memory and space information. Based on spike recordings of hippocampal neuronal populations, complex spiking patterns can be followed across extended periods including their modification of input-free self-generated patterns after environmental events (Buzsáki, 2004). Specific spiking sequences, which were elicited by experimental task design, have been shown to be re-enacted spontaneously during quiet wakefulness and sleep (Hartley et al., 2014; ONeill et al., 2010). Moreover, neuronal spike sequences measured in hippocampal place cells of rats featured re-occurrence directly after experimental trials as well as directly before (prediction of) upcoming experimental trials (Diba and Buzsáki, 2007).

Similar spiking patterns in hippocampal neurons during rest and sleep have been proposed to be critical in communicating local information to the neocortex for long-term storage, potentially including DMN regions. Moreover, in mice, invasively triggering spatial experience recall in the HC during sleep has been demonstrated to subsequently alter action choice during wakefulness (De Lavilläon et al., 2015). These HC-subserved mechanisms conceivably contribute to advanced cognitive processes that require re-experiencing or newly constructed mental scenarios, such as in recalling autobiographical memory episodes (Hassabis et al., 2007). Thus, the HC would orchestrate re-experience of environmental aspects for consolidations based on re-enactment and for integration into rich mental scene construction (Deuker et al., 2016; Bird et al., 2010). As such, the HC may impact ongoing perception of and action on the environment (Zeidman and Maguire, 2016; De Lavilléon et al., 2015).

### 2.4 The right and left TPJ: prediction error signaling and world semantics

The DMN emerges with its midline structures early in human development (Doria et al., 2010), while the right and left TPJs may become fully integrated into this major brain network only after birth. The TPJs are known to exhibit hemispheric differences based on microanatomical properties and cortical gyrification patterns (Seghier, 2013). Globally, neuroscientific investigations on hemispheric functional specialization have highlighted the right cerebral hemisphere as dominant for attentional functions and the left side for semantic functions (Seghier, 2013; Bzdok et al., 2013b, 2016a; Stephan et al., 2007).

The TPJ in the right hemisphere (RTPJ) has been shown to be closely related to multi-sensory prediction and prediction error signaling. This DMN region is probably central for action initiation during goal-directed psychological tasks and for sensorimotor behavior by integrating multi-sensory attention (Corbetta and Shulman, 2002). Its involvement was repeatedly reported in multi-step action execution (Hartmann et al., 2005), visuo-proprioceptive conflict (Balslev et al., 2005), and detection of environmental changes across visual, auditory, or tactile stimulation (Downar et al., 2000). Direct electrical stimulation of the human RTPJ during neurosurgery was associated with altered perception and stimulus awareness (Blanke et al., 2002). It was argued that the RTPJ encodes actions and predicted outcomes, without necessarily relating these neural processes to value estimation (Liljeholm et al., 2013; Hamilton and Grafton, 2008; Jakobs et al., 2009). Additionally, neural activity in the RTPJ has been proposed to reflect stimulus-driven attentional reallocation to self-relevant and unexpected sources of information as a circuit breaker that recalibrates functional control of brain networks (Bzdok et al., 2013b; Corbetta et al., 2008). Indeed, neurological patients with RTPJ damage have particular difficulties with multi-step actions (Hartmann et al., 2005). In the face of large discrepancies between actual and previously predicted environmental events, the RTPJ acts as a potential switch between externally-oriented mind sets focussed on the sensory environment and internally-oriented mind sets focussed on mental scene construction. For instance, temporally induced RTPJ damage in humans diminished the impact of predicted intentions of other individuals (Young et al., 2010), a capacity believed to be enabled by the DMN. The RTPJ might hence be an important relay that shifts away from the internally directed baseline processes to, instead, deal with unexpected environmental stimuli and events.

The left TPJ of the DMN (LTPJ), in turn, has a close relationship to Wernicke’s area involved in semantic processes, such as in spoken and written language. Neurological patients with damage in Wernicke’s area have a major impairment of language comprehension when listening to others or reading a book. Patient speech preserves natural rhythm and normal syntax, yet the voiced sentences lack meaning (i.e., aphasia). Abstracting from speech interpretations in linguistics and neuropsychology, the LTPJ appears to mediate access to and binding of world knowledge, such as required during action considerations (Binder and Desai, 2011; Seghier, 2013). Consistent with this view, LTPJ damage in humans also entailed problems in recognizing others’ pantomimed action towards objects without obvious relation to processing explicit language content (Varney and Damasio, 1987). Inner speech also hinges on knowledge recall about the physical and social world. Indeed, the internal production of verbalized thought (“language of the mind”) was closely related to the LTPJ in a pattern analysis of brain volume (Geva et al., 2011). Further, episodic memory recall and mental imagery to forecast future events strongly draw on re-assembling world knowledge. Isolated building blocks of world structure get rebuilt in internally constructed mental scenarios that guide present action choice, weigh hypothetical possibilities, and forecast event outcomes. Neural processes in the LTPJ may hence contribute to the automated predictions of the environment by incorporating experience-derived building blocks of world regularities into ongoing action, planning, and problem solving.

## 3 Reinforcement learning control: a process model for DMN function

We argue the outlined neurobiological properties of the DMN regions to be sufficient for implementing all components of a full-fledged reinforcement-learning (RL) system. Recalling past experience, considering candidate actions, random sampling of possible experiences, as well as estimation of instantaneous and expected delayed reward outcomes are key components of intelligent RL agents that are plausible to functionally intersect in the DMN.

RL is an area of machine learning concerned with searching optimal behavioral strategies through interactions with an *environment* with the goal to maximize some *cumulative reward.* The optimal behavior typically takes the future into account as some rewards could be *delayed.* Through repeated action on and feedback from the environment, the agent learns how to reach goals and continuously improve the collection of reward signals in a trial-and-error fashion (Fig. 3). At a given moment, each taken *action a* triggers a change in the *state* of the environment *s* → *s’*, accompanied by environmental feedback signals as *reward r = r(s,a,s’*) obtained by the agent. If the collected reward outcome yields a negative value it can be more naturally interpreted as *punishment.* In this setting, the environment is partially controlled by the action of the agent and the reward can be thought of as satisfaction or aversion that accompany the execution of a particular action.

**Fig. 3.**
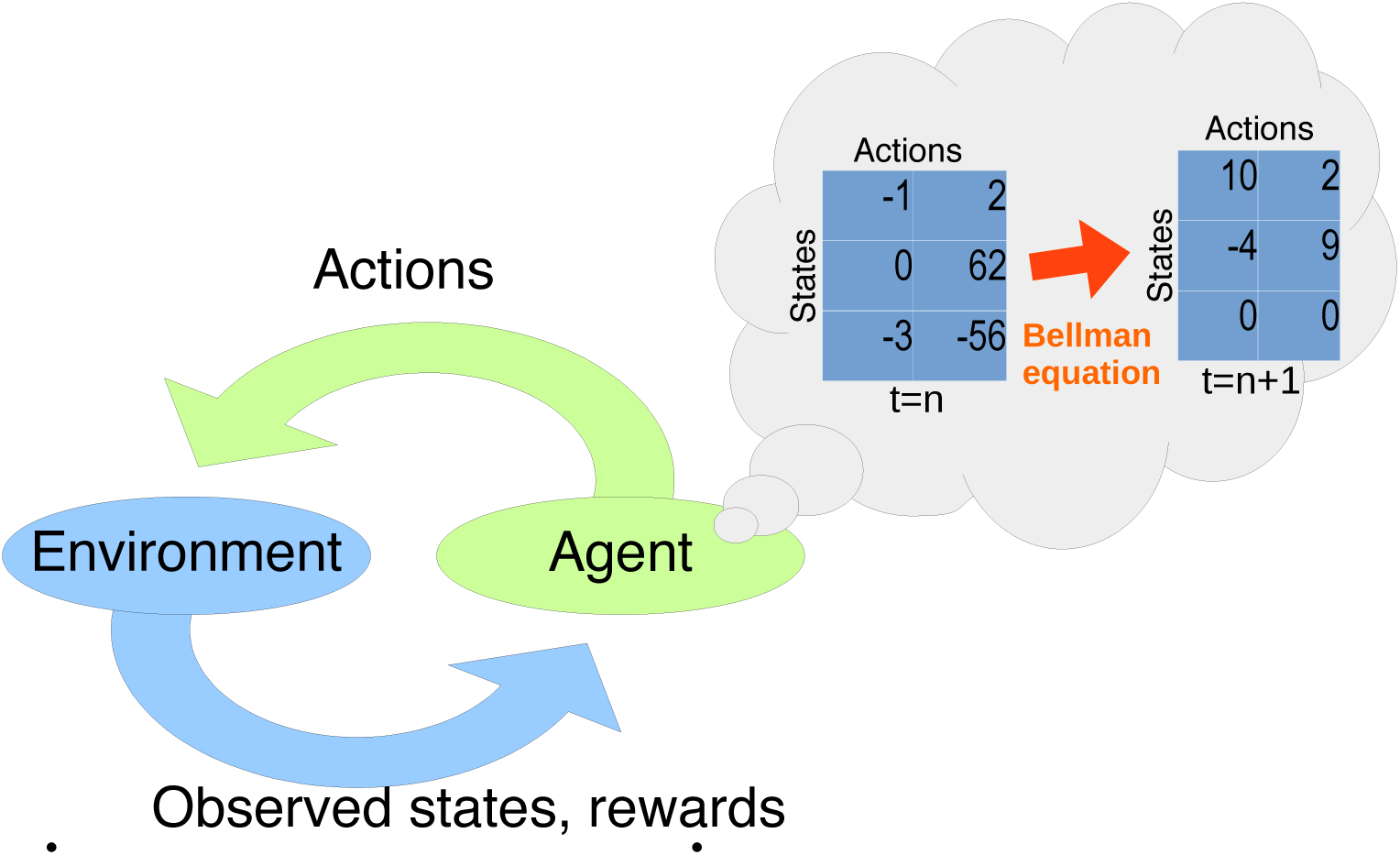
Reinforcement learning in a nutshell. Given the current state of the environment, the agent takes an action by following the policy matrix as updated by the Bellman equation. The agent receives a triggered reward and observes the next state. The process goes on until interrupted or a goal state is reached.

The environment is generally taken as *stochastic*, that is, changing in random ways. In addition, the environment is only *partially observable* in the sense that only limited aspects of the environment’s state are accessible to the agent’s sensory input (Starkweather et al., 2017). We assume that volatility of the environment is realistic in a computational model which sets out to explain DMN functions of the human brain. We argue that a functional account of the DMN based on RL can naturally embed human behavior in the tension between exploitative action with immediate gains and explorative action with longer-term reward outcomes (Dayan and Daw, 2008). In short, DMN implication in a diversity of particularly sophisticated human behaviors can be parsimoniously explained as instantiating probabilistic simulations of experience coupled with prediction error minimization to calibrate action trajectories for reward outcome maximization at different time-scales. Such a purposeful optimization objective may be subserved by a stochastic approximation based on a brain implementation of MCMC sampling.

### 3.1 Markov decision processes

RL has had considerable success in modeling many real-world problems, including super-human performance in complex video games (Mnih et al., 2015), robotics (Ng et al., 2004; Abbeel and Ng, 2004), and strategic board games like the breakthrough results upon recently on the game of Go (Silver et al., 2016) considered to be a milestone benchmark in artificial intelligence. In artificial intelligence and machine learning, a popular computational model for multi-step decision processes in such an environment are MDPs (Sutton and Barto, 1998). An MDP operationalizes a sequential decision process in which it is assumed that environment dynamics are determined by a Markov process, but the agent cannot directly observe the underlying state. Instead, the agent tries to optimize a *subjective* reward signal (i.e., likely to be different for another agent in the same state), by maintaining probability distributions over actions according to their expected utility. This is a minimal set of assumptions that can be made about an environment faced by an agent engaged in interactive learning.

#### Definition.

*Mathematically, an MDP is simply a quintuplet* (*𝒮, 𝒜,r,p*) where

- *𝒮 is the set of states, such as 𝒮 = {happy, sad, puzzeled}.*
- *𝒜 is the set of actions, such as 𝒜 = {read, run, laugh, sympathize, empathize}.*
- *r:𝒮 × 𝒜 × 𝒮 → is the reward function, so that r(s, a, s’) is the instant reward for taking action a in state s followed by a state-transition s* → *s’.*
- *p*: *𝒮* × *𝒜* × *𝒮* → [0,1], (*s,a,s’*)↦*p*(*s*’|*s,a*), the probability of moving to state s’ if action a is taken from state *s*. In addition, one requires that such transitions be Markovian. Consequently, the future states are independent of past states and only depend on the present state and action taken.

The process has *memory* if the subsequent state depends not only on the current state but also on a number of past states. Rational probabilistic planning can thus be reformulated as a standard memoryless Markov process by simply expanding the definition of the state *s* to include experience episodes of the past. This extension adds the capacity for memory to the model because the next state then depends not only on the current situation but also on previously experienced events, which is the motivation behind Partially Observable MDPs (POMDPs) (Starkweather et al., 2017; O’Reilly and Frank, 2006). Nevertheless, this mathematical property of POMDPs mostly accounts for implicit *memory.* Since the current paper is concerned with plausibility at the behavioral and neurobiological level, we will address below how our account can accommodate the neurophysiological constraints of the DMN and the explicit *memory* characteristics of human agents.

**Why Markov Decision Processes?**

One may wonder whether MDP models are applicable to something as complex as human behavior. For instance, financial trading is largely a manifestation of strategic decision-making of interacting human agents. According to how the market responds, the agent incurs gain or loss as environmental feedback of the executed financial actions. Recent research on automatizing market exchanges by algorithmic trading has successfully used MDPs as a framework for modeling these elaborate behavioral dynamics (Brazdil et al., 2017; Yang et al., 2015, 2014, 2012; Dempster and Leemans, 2006;

Hult and Kiessling, 2010; Abergel et al., 2017). MDPs have also been effective as a behavioral model in robotics (Ng et al., 2004; Abbeel and Ng, 2004) and in challenging multistep strategy games (Mnih et al., 2015; Silver et al., 2016; Pritzel et al., 2017). As such, we aim to expand MDP applications as a useful model from “online” decision-making to the realms of “offline” behaviors most associated with the DMN.

**Towards model-free reinforcement learning for the DMN.**

Model-free RL can be plausibly realized in the human brain (O’Doherty et al., 2015; Daw and Dayan, 2014). Indeed, it has been proposed (Gershman et al., 2015) that a core property of human intelligence is the improvement of expected utility outcomes as a strategy for action choice in uncertain environments, a situation perfectly captured by the formalism of MDPs. It has also long been proposed (Dayan and Daw, 2008) that there can be a direct mapping between model-free RL learning algorithms and aspects of the brain. The neurotransmitter dopamine could serve as a teaching signal’ to better estimate value associations and action policies by controlling synaptic plasticity in the reward-processing circuitry, including the NAc. In contrast, *model-based RL* would start off with some mechanistic assumptions about the dynamics of the world. These can be assumptions about the physical laws governing the agent’s environment or constraints on the state space, transition probabilities between states, reward contingencies, etc. An agent might represent such knowledge about the world as follows:

- *r*(*s, “stand still”*) = 0 if *s* does not correspond to a location offering relevant resources.
- *p*(*s’|s, “stand still”*) = 1 if *s’* = *s* and 0 otherwise.
- etc.

Such knowledge can be partly extracted from the environment: the agent infers a model of the world while learning to take optimal decisions based on the current representation of the environment. These methods learn what the effect is going to be of taking a particular action in a particular state. The result is an estimate of the underlying MDP which can then be either solved exactly or approximately, depending on the setting and what is feasible.

In contrast, model-free methods require no prespecified knowledge of the environment (transition probabilities, types of sensory input, etc.) or representation thereof. The agent infers which state-action pairs lead to reward through sampling the world in a trial-and-error manner and derives longer-term reward aggregates using environmental feedback information as an incentive. In so doing, model-free agents ultimately learn both an action policy and an implicit representations of the world. This distinction between model-free and model-based RL is similar to previous views (Dayan and Berridge, 2014).

#### 3.1.1 Accumulated rewards and policies

The behavior of the agent is governed by a *policy*, which maps states of the world to probability distributions over candidate actions. Starting at time *t* = 0, following a policy *π* generates a trajectory of action choices:

> **choose action:** *a*_0_ ~ *π*(*a*|*s*_0_)
>
> **observe transition:** *s*_1_ ~ *p*(*s*|*s*_0_,*a*_0_) **and collect reward** *R*_1_ = *r*(*s*_1_,*a*_1_, *s*_2_)
>
> **choose action:** *a*_1_ ~ *π*(*a*|*s*_*t*_)
>
> **observe transition:** *s*_2_~*p*(*s*|*s*_1_,*a*_1_),**and collect reward** *R*_1_ = *r*(*s*_1_,*a*_1_, *s*_2_)
>
> ⋮
>
> **choose action:** *a*_*t*_ ~*π*(*a* |*s*_*t*_)
>
> **observe transition:***s*_*t*+1_ ~*p*(*s* |*s*_*t*_,*a*_*t*_),**and collect reward** *R*_*t*_ = *r*(*s*_*t*_,*a*_*t*_,*s*_*t*+1_)
>
> ⋮

We assume time-invariance in that we expect the dynamics of the process to be equivalent over sufficiently long time windows of equal length (i.e., stationarity). Since an action executed in the present moment might have repercussions in the far future, it turns out that the quantity to optimize is not the instantaneous rewards *r(s,a*), but a *cumulative reward* estimate which takes into account expected reward from action choices in the future. A common approach to modeling this gathered outcome is the time-discounted cumulative reward

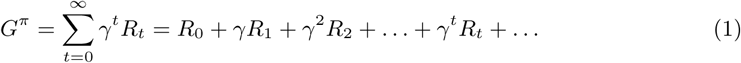

This random variable measures the cumulative reward of following an action policy *π*.Note that value buffering may be realized in the vmPFC. This DMN region has direct connections to to the NAc, known to be involved in reward evaluation.

The goal of the RL agent is then to successively update this action policy in order to maximize G^*π*^ on average (cf. below). In (1), the definition of cumulative reward *G*^*π*^, the constant *γ* (0 ≤ *γ* < 1) is the *reward discount factor*, viewed to be characteristic for a certain agent. On the one hand, setting *γ* = 0 yields perfectly hedonistic behavior. An agent with such a shortsighted time horizon is exclusively concerned with immediate rewards. This is however not compatible with coordinated planning of longer-term agendas that is potentially subserved by neural activity in the DMN. On the other hand, setting 0 < *γ* < 1 allows a learning process to arise. A positive *γ* can be seen as calibrating risk-seeking trait of the intelligent agent, that is, the behavioral predispositions related to trading longer delays for higher reward outcomes. Such an agent puts relatively more emphasis on rewards expected in a more distant future. Concretely, rewards that are not expected to come within *τ* ≔ 1/(1 — *γ*) time steps from the present point are ignored. The complexity reduction by time discounting alleviates the variance of expected rewards accumulated across considered action cascades by limiting the depth of the search tree. Given that there is more uncertainty in the far future, it is important to appreciate that a stochastic policy estimation is more advantageous in many RL settings.

### 3.2 The components of reinforcement learning in the DMN

Given only the limited information available from an MDP, at a state *s* the average utility of choosing an action *a* under a policy *π* can be captured by the single number

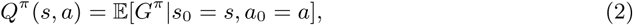

called the *Q*-value for the state-action pair (*s,a*). In other words, *Q*^*π*^(*s,a*) corresponds to the expected reward over all considered action trajectories, in which the agent sets out in the environment in state *s*, chooses action *a*, and then follows the policy *π* to select future actions. For the brain, *Q*^*π*^(*s,a*) defined in (2) provides the subjective utility of executing a specific action. It thus answers the question “What is the expected utility of choosing action *a*, and its ramifications, in this situation?”. *Q*^*π*^(*s,a*) offers a formalization of optimal behavior that may well capture such processing aspects subserved by the DMN in human agents.

#### 3.2.1 Optimal behavior and the Bellman equation

Optimal behavior of the agent corresponds to a strategy *π*^*^ for choosing actions such that, for every state, the chosen action guarantees the best possible reward on average. Formally,

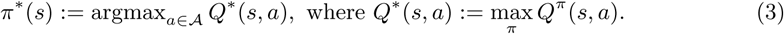

The learning goal is to approach the policy *π*\* as close as possible, that is to solve the MDP. Note that (3) presents merely a definition and does not lend itself as a candidate schema for solving MDPs with even moderately sized action and state spaces (i.e., intractability). Fortunately, the *Bellman equation* (Sutton and Barto, 1998) provides a fixed-point relation which defines *Q*^*^ implicitly via a sampling procedure, without querying the entire space of policies, with the form

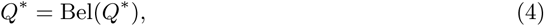

Random as it depends both on the environment’s dynamics and the policy *π* being executed. The exponential delay discounting function used here refers to the usual formulation in the field of reinforcement learning, although psychological experiments may also reveal other discounting regimes (Green and Myerson, 2004).
where the so-called Bellman transform Bel(Q) of an arbitrary *Q*-value function *Q*: *S* x *A* →R is another *Q*-value function defined by

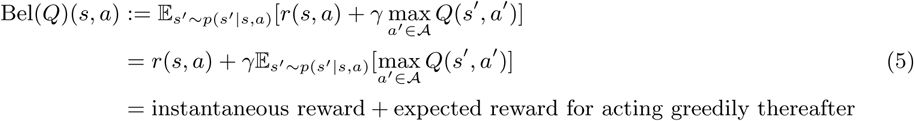

The Bellman equation (4) is a temporal consistency equation which provides a dynamic decomposition of optimal behavior by dividing the *Q* value function into the immediate reward and the discounted rewards of the upcoming states. The optimal Q-value operator *Q*\* is a fixed point for this equation. As a consequence of this outcome stratification, the complicated dynamic programming problem (3) is broken down into simpler sub-problems at different time points. Indeed, exploitation of hierarchical structure in action considerations has previously been related to the medial prefrontal part of the DMN (Koechlin et al., 1999; Braver and Bongiolatti, 2002). Using the Bellman equation, each state can be associated with a certain value to guide action towards a preferred state, thus improving on the current action policy of the agent. Note that in (4) the random sampling is performed only over quantities which depend on the environment. This aspect of the learning process can unroll off-policy by observing state transitions triggered by another (possibly stochastic) behavioral policy.

##### Box 1: Neural correlates of the Bellman equation in the DMN

Relating decomposition of consecutive action choices by the Bellman equation to neuroscientific insights, specific neural activity in the dorsal prefrontal cortex (BA9) was linked to processing “goal-tree sequences” in human brain-imaging experiments (Koechlin et al., 1999, 2000). Sub-goal exploration may require multi-task switching between cognitive processes as later parts of a solution frequently depend on respective earlier steps in a given solution path, which necessitates storage of expected intermediate outcomes. As such, “cognitive branching” operations for nested processing of behavioral strategies are likely to entail secondary reallocation of attention and working-memory resources. Further brain-imaging experiments corroborated the prefrontal DMN to subserve “processes related to the management and monitoring of sub-goals while maintaining information in working memory” (Braver and Bongiolatti, 2002) and to functionally couple with the hippocampus conditioned by “deep versus shallow planning” (Kaplan et al., 2017). Moreover, neurological patients with lesions in this DMN region were reported to be impaired in aspects of realizing “multiple sub-goal scheduling” (Burgess et al., 2000). Hence, the various advanced human abilities subserved by the DMN, such as planning and abstract reasoning, can be viewed to involve some form of action-decision branching to enable higher-order executive control.

#### 3.2.2 Value approximation and the policy matrix

As already mentioned in the previous section, Q-learning (Watkins and Dayan, 1992) optimizes over the class of deterministic policies of the form (3). State spaces may be extremely large and tracking all possible states and actions may require prohibitively excessive computation and memory resources. The need of maintaining an explicit table of states can be eliminated by instead using of an approximate *Q*-value function 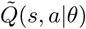 by keeping track of an approximating parameter *θ* of much lower dimension than the number of states. At a given time step, the world is in a state *s* ∈ *S*, and the agent takes an action which it expects to be the most valuable on average, namely

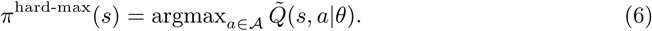

This defines a mapping from states directly to actions. For instance, a simple linear model with a kernel *ϕ* would be of the form 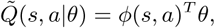where *ϕ*(*s,a*) would represent a high-level representation of the state-action pairs (*s,a*), as was previously proposed (Song et al., 2016), or artificial neural-network models as demonstrated in recent seminal investigations (Mnih et al., 2015; Silver et al., 2016) for playing complex games (atari, Go, etc.) at super-human levels. In the DMN, the dmPFC would implement such a hard-max lookup over the action space. The model parameters *θ* would correspond to synaptic weights and connection strengths within and between brain regions. It is a time-varying neuronal program which dictates how to move from world states *s* to actions *a* via the hard-max policy (6). The approximating *Q*-value function 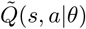 would inform the DMN with the (expected) usefulness of choosing an action *a* in state *s.* The DMN, and in particular its dmPFC part, could then contribute to the choice, at a given state *s*, of an action *a* which maximizes the approximate *Q*-values. This mapping from states to actions that is conventionally called *policy matrix* (Mnih et al., 2015; Silver et al., 2016). Learning consists in starting from a given table and updating it during action choices, which take the agent to different table entries.

#### 3.2.3 Self-training and the loss function

Successful learning in brains and computer algorithms may not be possible without a defined learning goal the *loss function.* The action *a* chosen in state *s* according to the policy matrix defined in (6) yields a reward *r* collected by the agent, after which the environment transitions to a new state *s*’ *∈ S.* One such cycle yields a new *experience e* = (*s,a,r,s’*). Each cycle represents a behavior unit of the agent and is recorded in replay memory buffer which we hypothesize to be subserved by the HC —, possibly discarding the oldest entries to make space: *D*← append(*D, e*). At time step *k*, the agent seeks an update *θ*_*k*_ ← *θ*_*k*-1_ + *δθ*_*k*_ of the parameters for its approximate model of the Q-value function. This warrants a learning process and definition of a loss function. The Bellman equation (4) provides a way to obtain such a loss function (9) as we outline in the following. Experience replay consists in sampling batches of experiences *e* (*s, a, r, s’*) ~ *D* from the replay memory *D*. The agent then tries to approximate the would-be Q-value for the state-action pair (*s,a*) as predicted by the Bellman equation (4), namely

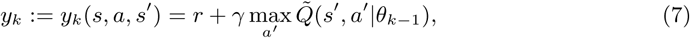

with the estimation of a parametrized regression model 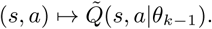From a neurobiological perspective, experience replay can be manifested as the re-occurrence of neuron spiking sequences that have also been measured during specific prior actions and environmental states. The HC is a strong candidate for contributing to such neural reinstantiation of behavioral episodes as neuroscience experiments have repeatedly indicated in rats, mice, cats, rabbits, songbirds, and monkeys (Buhry et al., 2011; Nokia et al., 2010; Dave and Margoliash, 2000; Skaggs et al., 2007).

At the current step k, computing an optimal parameter update then corresponds to finding the model parameters *θ*_*k*_ which minimize the following mean-squared loss function

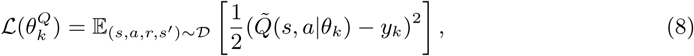

where *yk* is obtained from (4). A recently proposed, practically successful alternative approach is to learn this representation using an artificial deep neural-network model. This approach leads to the so-called *deep Q-learning* (Mnih et al., 2015; Silver et al., 2016) family of methods which is the current state-of-the-art in RL research. The set of model parameters *θ* that instantiate the non-linear interactions between layers of the artificial neural network may find a neurobiological correspondence in the adaptive strengths of axonal connections between neurons from the different levels of the neural processing hierarchy (Mesulam, 1998; Taylor et al., 2015).

**A note on bias in self-training.** Some bias may be introduced by self-training due to information shortage caused by the absence of external stimulation. One way to address this issue is using importance sampling to replay especially those state-transitions from which there is more to learn for the agent (Schaul et al., 2015; Hessel et al., 2017). New transitions are inserted into the replay buffer with maximum priority, thus shifting emphasis to more recent transitions. Such insertion strategy would help counterbalance the bias introduced by the information shortage incurred by absent external input. Other authors noticed (Hessel et al. 2017) that such prioritized replay reduces the data complexity and the agent shows faster increases in learning performance.

#### 3.2.4 Optimal control via stochastic gradient descent in the DMN

Efficient learning of the entire set of model parameters can effectively be achieved via stochastic *gradient descent*, a universal algorithm for finding local minima based on the first derivative of the optimization objective. Stochastic here means that the gradient is estimated from batches of training samples, which here corresponds to blocks of experience from the replay memory:

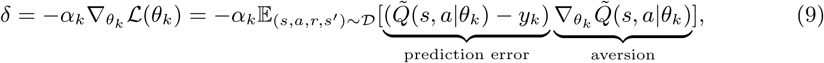

where the positive constants *α*_1_, *α*_2_,… are learning rates. Thus, the subsequent action is taken to drive reward prediction errors to percolate from lower to higher processing layers to modulate the choice of future actions. It is known that under special conditions on the learning rates *α*_*k*_ namely that the learning rates are neither too large nor too small, or more precisely that the sum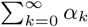 diverges while 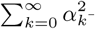the thus generated approximating sequence of Q-value functions

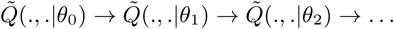

are attracted and absorbed by the optimal *Q*-value function *Q*\* defined implicitly by the Bellman equation (4).

#### 3.2.5 Does the hippocampus subserve MCMC sampling?

In RL, MCMC simulation is a common means to update the agent’s belief state based on stochastic sampling around states and possible transitions (Daw and Dayan, 2014). MCMC simulation provides a simple method for evaluating the value of a state. This inference procedure provides an effective mechanism both for tree search (of the considered action trajectories) and for belief state updates, breaking the curse of dimensionality and allowing much greater scalability than an RL agent without stochastic resampling procedures. Such methods have scaling as a function of available data (i.e., sample complexity) that is determined only by the underlying difficulty of the MDP, rather than the size of the state space or observation space, which can be prohibitively large.

In the human brain, the HC could contribute to synthesizing imagined sequences of world states, actions and rewards (Aronov et al., 2017; Chao et al., 2017; Boyer, 2008). These simulations of experience batches would be used to update the value function, without ever looking inside the black box describing the model’s dynamics. A brain-imaging experiment in humans for instance identified hippocampal signals that specifically preceded upcoming choice performance in prospective planning in new environments (Kaplan et al., 2017). This would be a simple control algorithm by evaluating all legal actions and selecting the action with highest expected cumulative rewards. In MDPs, MCMC simulation provides an effective mechanism both for tree search and for belief-based state updates, breaking the curse of dimensionality and allowing much greater scalability than has previously been possible (Silver et al., 2016). This is because expected consequences of action choices can be well evaluated although only a subset of the states are actually considered (Daw and Dayan, 2014).

**A note on implicit and explicit memory.**While Markov processes are usually memoryless, it is mathematically feasible to incorporate the previous states of such model into the current state. This extension may partially account for implicit memory at the behavioral level, but may not explain the underlying neurobiological implementation or accommodate explicit memory. Implicit memory-based processing arises in our MDP account of DMN function in several different forms: successive updates of a) the action policy and the value function, both being products of the past, as well as b) the deep non-linear relationships within the hierarchical connections of biological neural networks. The brain’s adaptive synaptic connections can be thought as a deep neural-network architecture affording an implicit form of information compression of life experience. Such memory traces are stored in the neural machinery and can be implicitly retrieved as a form of knowledge during simulation of action rather than accessed as a stored explicit representation (Pezzulo, 2011). c) Certain neural processes in the hippocampus can be seen as some type of MCMC sampling for memory recall, which can also be a basis for probabilistic simulations across time-scales (Schacter et al., 2007; Axelrod et al., 2017).

### 3.3 Putting everything together

The DMN is today known to consistently increase in neural activity when humans engage in cognitive processes that are relatively detached from the current sensory environment. The more familiar and predictable the current environment, the more brain resources may remain for allocating DMN activity to MDP processes extending beyond the present time and sensory context. In line with this perspective, DMN engagement was shown to heighten and relate to effective behavioral responses in the practiced phase of a demanding cognitive flexibility task, as compared to acquisition phase when participants learned context-specific rules. This involvement in automated decision-making has led the author
s to propose an “autopilot” role for the DMN (Vatansever et al., 2017), which may contribute to optimizing control of the organism in general. Among all parts of the DMN, the RTPJ is perhaps the most evident candidate for a network-switching relay that calibrates between processing of environment-engaged versus internally generated information (Downar et al., 2000; Golland et al., 2006; Bzdok et al., 2013b).

Additionally, the DMN was proposed to be situated at the top of the brain network hierarchy, with the subordinate salience and dorsal attention network in the middle and the primary sensory cortices at the bottom (Carhart-Harris and Friston, 2010; Margulies et al., 2016b). Its putative involvement in thinking about hypothetical experiences and future outcomes appears to tie in with the implicit computation of action and state cascades as a function of experienced events and collected feedback from the past. A policy matrix encapsulates the choice probabilities of possible actions on the world given a current situation (i.e., state). The DMN may subserve constant exploration of candidate action trajectories and nested estimaton of their cumulative reward outcomes. Implicit computation of future choices provides a potential explanation for the evolutionary emergence and practical usefulness of mind-wandering at day-time and dreams during sleep in humans.

**Fig. 4.**
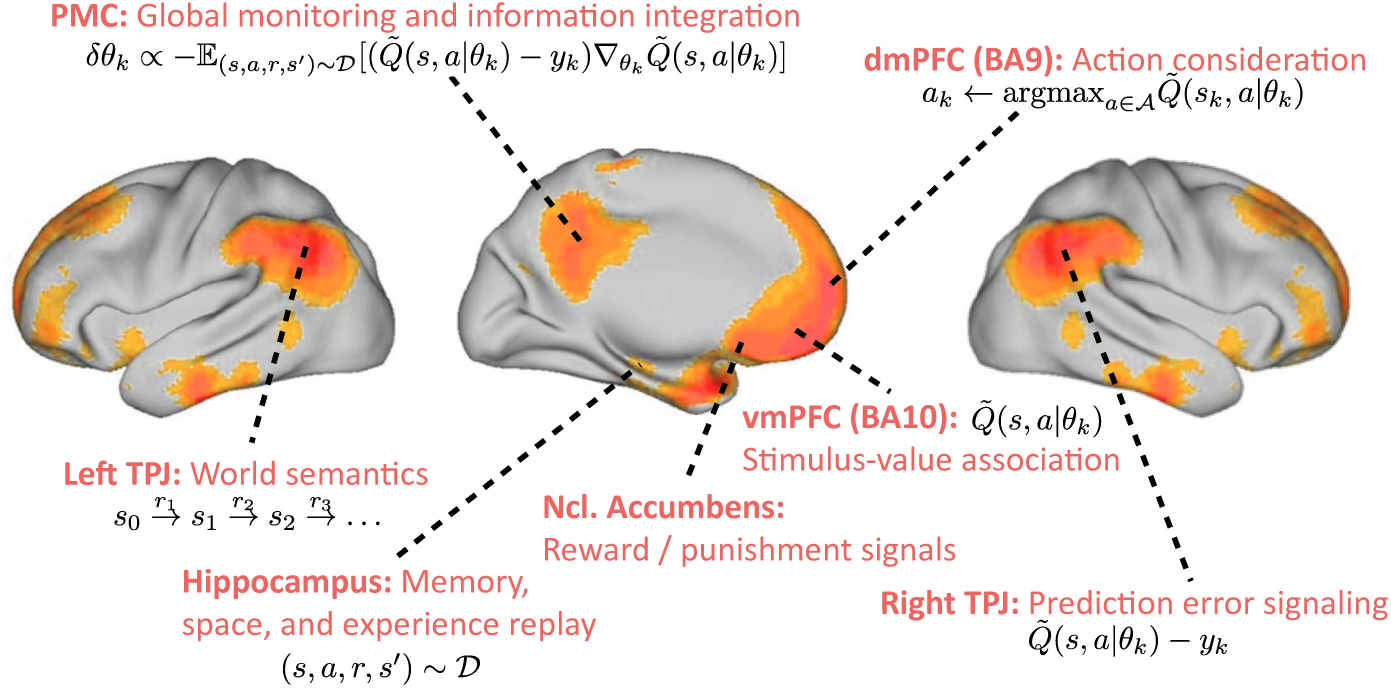
Default mode network: possible neurobiological implementation of reinforcement learning. Overview of how the constituent regions of the DMN (refer to section 2) may map onto computational components necessary for an RL agent.

The HC may contribute to generating perturbed action-transition-state-reward samples as batches of pseudo-experience (i.e., recalled, hypothesized, and forecasted scenarios). The small variations in these experience samplings allow searching a larger space of model parameters and possible experiences. Taken to its extreme, stochastic recombination of experience building blocks can further optimize the behavior of the RL agent by model learning from scenarios in the environment that the agent might only very rarely or never encounter. An explanation is thus offered for experiencing seemingly familiar situations that a human has however never actually encountered (i.e., déja vu effect). While such a situation may not have been experienced in the physical world, the DMN may have previously stochastically generated, evaluated, and adapted to such a randomly synthesized event. Generated representations arguably are “internally manipulable, and can be used for attempting actions internally, before or instead of acting in the external reality, and in diverse goal and sensory contexts, i.e. even outside the context in which they were learned” (Pezzulo, 2011). In the context of scarce environmental input and feedback (e.g., mind-wandering or sleep), mental scene construction allows pseudo-experiencing possible future scenarios and action outcomes.

From the perspective of a model-free RL agent, prediction in the DMN reduces to generalization of policy and value computations from sampled experiences to successful action choices and reward predictions in future states. As such, plasticity in the DMN arises naturally. If an agent behaving optimally in a certain environment moves to new, yet unexperienced environment, reward prediction errors will largely increase. This feedback will lead to adaptation of policy considerations and value estimations until the intelligent system converges to a new steady state of optimal action decisions in a volatile world.

#### Box 2: Proposed studies for testing the MDP account of DMN function

1. Experiment (Humans): We hypothesize a functional relationship between the DMN closely associated with the occurrence of stimulus-independent thoughts and the reward circuitry. During an iterative neuroeconomic two-player game, fMRI signals in the DMN could be used to predict reward-related signals in the nucleus accumbens across trials in a continuous learning paradigm. We expect that the more DMN activity is measured to be increased, supposedly the higher the tendency for stimulus-independent thoughts, the more the fMRI signals in the reward circuits should be independent of the reward context in the current sensory environment.

2. Experiment (Humans): We hypothesize a functional dissociation between computations pertaining to action policy versus adapting stimulus-value associations as we expect implementation in different subsystems of the DMN. First, we expect that fMRI signals in the right temporo-parietal junction relate to behavioral changes subsequent to adaptation in the action choice tendencies (policy matrix) involved in non-value-related prediction error. Second, fMRI signals in the ventromedial prefrontal cortex should relate to behavioral changes following adaptation in value estimation (value matrix) due to reward-related stimulus-value association. We finally expect that fMRI signals in the posteromedial cortex, as a potential global information integrator, are related to shifts in overt behavior based on previous adaptations in both policy or value estimation.

3. Experiment (Animals): We hypothesize that experience replay for browsing problem solutions subserved by the DMN is contributing to choice behavior in mice. Hippocampal single-cell recordings have shown that neural patterns during experimental choice behavior are reiterated during sleep and before making analogous choices in the future. Necessity of cortical DMN regions, in addition to the hippocampus, for mind-searching candidate actions during choice behavior can be experimentally corroborated by causal disruption of DMN regions, such as by circumscribed brain lesion or optogenetic intervention in the inferior parietal and prefrontal cortices.

4. Experiment (Humans): We hypothesize that the relevant time horizon is modulated by various factors such as age, acute stress and time-enduring impulsivity traits. Using a temporal discounting experiment, it can be quantified how the time horizon is affected at the behavioral level and then traced-back to its corresponding neural representation. Such experimental investigation can be designed to examine between-group and within-group effects (e.g., impulsive population like chronic gamblers or drug addicts); and brought in context with the participants age and personality traits.

5. Experiment (Humans & Animals): An additional layer of learning concerns the addition of new entries in the state and action spaces. Extension of the action repertoire could be biologically realized by synaptic epigenesis (Gisiger et al., 2005). Indeed, the tuning of synaptic weights through learning can stabilize additional patterns of activity by creating new attractors in the neural dynamics landscape (Takeuchi et al., 2014). Those attractors can then constrain both the number of factors taken into account by decision processes and the possible behaviors of the agent (Wang, 2008). To examine this potential higher-level mechanism, we propose to probe how synaptic epigenesis is related to neural correlates underlying policy matrix updates: in humans the changes of functional connectivity between DMN regions can be investigated following a temporal discounting experiment and in monkeys or rodents anterograde tracing can be used to study how homolog regions of the DMN present increased synaptic changes compare to other parts of the brain.

**Fig. 5.**
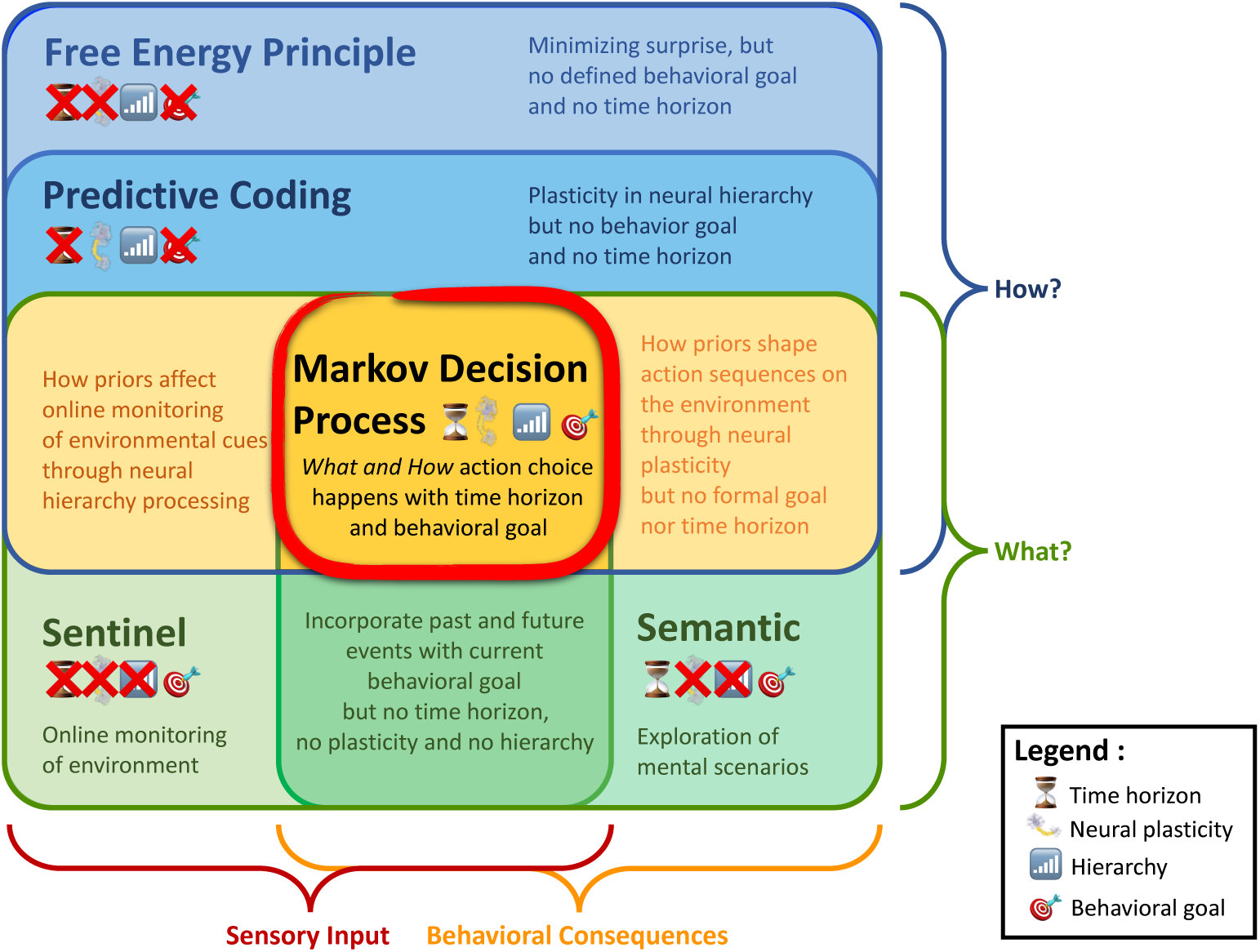
Situating Markov Decision Processes among other accounts of default mode function. The Venn diagram summarizes the relationship between four previously proposed explanations for the functional role of the DMN and our present account. Viewing empirical findings in the DMN from the MDP viewpoint incorporates important aspects of the free energy principle, predictive coding, sentinal hypothesis, and semantic hypothesis. The MDP account may reconcile several strengths of these functional accounts in a process model that simultaneously acknowledges environmental input and behavioral choices as well as the computational and algorithmic properties (How? and What?) underlying higher-order control of the organism.

## 4 Relation to existing accounts

### 4.1 Predictive coding hypothesis

Predictive coding mechanisms (Clark, 2013; Friston, 2008) are a frequently evoked idea in the context of default mode function (Bar et al., 2007). Cortical responses are explained as emerging from continuous functional interaction between higher and lower levels of the neural processing hierarchy. Feed-forward sensory processing is constantly calibrated by top-down modulation from more multi-sensory and associative brain regions further away from primary sensory cortical regions. The dynamic interplay between cortical processing levels may enable learning about aspects of the world by reconciling gaps between fresh sensory input and predictions computed based on stored prior information. At each stage of neural processing, an internally generated expectation of aspects of environmental sensations is directly compared against the actual environmental input. A prediction error at one of the processing levels induces plasticity changes of neuronal projections to allow for gradually improved future prediction of the environment. In this way, the predictive coding hypothesis offers explanations for the constructive, non-deterministic nature of sensory perception (Friston, 2010; Buzsáki, 2006) and the intimate relation of motor movement to sensory expectations (Wolpert et al., 1995; Kárding and Wolpert, 2004). Contextual integration of sensorimotor perception-action cycles may be maintained by top-down modulation using internally generated information about the environment.

In short, predictive coding processes conceptualize updates of the internal representation of the environment to best accommodate and prepare the organism for processing the constant influx of sensory stimuli and performing action on the environment. There are hence a number of common properties between the predictive coding account and the proposed formal account of DMN function based on MDPs. Importantly, a generative model of how perceived sensory cues arise in the world would be incorporated into the current neuronal wiring. Further, both functional accounts are supported by neuroscientific evidence that suggest the human brain to be a “statistical organ” (Friston et al., 2014) with the biological purpose to generalize from the past to new experiences. Neuroanatomically, axonal back projections indeed outnumber by far the axonal connections mediating feedforward input processing in the monkey brain and probably also in humans (Salin and Bullier, 1995). These many and diverse top-down modulations from higher onto downstream cortical areas can inject prior knowledge at every stage of processing environmental information. Moreover, both accounts provide a parsimonious explanation for why the human brain’s processing load devoted to incoming information decreases when the environment becomes predictable. This is because the internal generative model only requires updates after discrepancies have occurred between environmental reality and its internally reinstantiated representation. Increased computation resources are however allocated when unknown stimuli or unexpected events are encountered by the organism. The predictive coding and MDP account hence naturally evoke a mechanism of brain plasticity in that neuronal wiring gets increasingly adapted when faced by unanticipated environmental challenges.

While sensory experience is a constructive process from both views, the predictive coding account frames sensory perception of the external world as a generative experience due to the modulatory top-down influence at various stages of sensory input processing. This generative top-down design is replaced in our MDP view of the DMN by a sequential decision-making framework. Further, the hierarchical processing aspect from predictive coding is re-expressed in our account in the form of nested prediction of probable upcoming actions, states, and outcomes. While both accounts capture the consequences of action, the predictive coding account is typically explained without explicit parameterization of the agent’s time horizon and has a tendency to be presented as emphasizing prediction about the immediate future. In the present account, the horizon of that look into the future is made explicit in the *γ* parameter of the Bellman equation. Finally, the process of adapting the neuronal connections for improved top-down modulation takes the concrete form of stochastic gradient computation and back-propagation in our MDP implementation. It is however important to note that the neurobiological plausibility of the back-propagation procedure is controversial (Goodfellow et al., 2016).

In sum, recasting DMN function in terms of MDPs therefore naturally incorporates a majority of aspects from the prediction coding hypothesis. The present MDP account of DMN function may therefore serve as a concrete implementation of predictive coding ideas. MDPs have the advantage of exposing an explicit mechanisms for controlling the horizon of future considerations and for how the internal representation of the world is updated, as well as why certain predictions may be more relevant to the agent than others.

### 4.2 Semantic hypothesis

This frequently proposed cognitive account to explain DMN function revolves around forming logical associations and abstract analogies between experiences and conceptual knowledge derived from past behavior (Bar, 2007; Binder et al., 1999; Constantinescu et al., 2016). Analogies might naturally tie incoming new sensory stimuli to explicit world knowledge (i.e., semantics) (Bar, 2009). The encoding of complex environmental features could thus be facilitated by association to known similar states. Going beyond isolated meaning and concepts extracted from the world, semantic building blocks may need to get recombined to enable mental imagery to (fore)see never-experienced scenarios. As such, semantic knowledge would be an important ingredient for optimizing behavior by constantly simulating possible future scenarios (Boyer, 2008; Binder and Desai, 2011). Such cognitive processes can afford the internal construction and elaboration of necessary information that is not presented in the immediate sensory environment by recombining building blocks of concept knowledge and episodic memories (Hassabis and Maguire, 2009). Indeed, in aging humans, remembering the past and imagining the future equally decreased in the level of detail and were associated with concurrent deficits in forming and integrating relationships between items (Addis et al., 2008; Spreng and Levine, 2006). Further, episodic memory, language, problem solving, planning, estimating others’ thoughts, and spatial navigation represent neural processes that are likely to build on abstract world knowledge and logical associations for integrating the constituent elements in rich and coherent mental scenes (Schacter et al., 2007). “[Foresight] and simulations are not only automatically elicited by external events but can be endogenously generated when needed. […] The mechanism of access via simulation could be a widespread method for accessing and producing knowledge, and represents a valid alternative to the traditional idea of storage and retrieval” (Pezzulo, 2011). Such mental scene-construction processes could contribute to interpreting the present and foreseeing the future. Further, mental scene imagery has been proposed to imply a distinction between engagement in the sensory environment and internally generated mind-wandering (Buckner and Carroll, 2007). These investigators stated that “A computational model […] will probably require a form of regulation by which perception of the current world is suppressed while simulation of possible alternatives are constructed, followed by a return to perception of the present”.

In comparison, both the semantic hypothesis and the present formal account based on MDPs expose mechanisms of how action considerations could be explored. In both accounts, there is also little reason to assume that contemplating alternative realities of various levels of complexity, abstraction, time-scale, and purpose rely on mechanisms that are qualitatively different. This interpretation concurs with DMN activity increases across time, space, and content domains demonstrated in many brain-imaging studies (Spreng et al., 2009; Laird et al., 2009; Bzdok et al., 2012; Binder et al., 2009). Further, the semantic hypothesis and MDP account offer explanations why HC damage does not only impair recalling past events, but also imagining hypothetical and future scenarios (Hassabis et al., 2007). While both semantic hypothesis and our formal account propose memory-enabled, internally generated information for probabilistic representation of action outcomes, MDPs render explicit the grounds on which an action is eventually chosen, namely, the estimated cumulative reward. In contrast to many versions of the semantic hypothesis, the MDPs naturally integrate the egocentric view (more related to current action, state, and reward) and the world view (more related to past and future actions, states, and rewards) on the world in a same optimization problem. Finally, the semantic account of DMN function does not provide suffcient explanation of how explicit world knowledge and logical analogies thereof lead to foresight of future actions and states. The semantic hypothesis does also not fully explain why memory recall for scene construction in humans is typically fragmentary and noisy instead of accurate and reliable. In contrast to existing accounts on semantics and mental scene construction, the random and creative aspects of DMN function are explained in MDPs by the advantages of stochastic optimization. Our MDP account provides an algorithmic explanation in that stochasticity of the parameter space exploration by MCMC approximation achieves better fine-tuning of the action policies and inference of expected reward outcomes. That is, the purposeful stochasticity of policy and value updates in MDPs provides a candidate explanation for why humans have evolved imperfect noisy memories as the more advantageous adaptation. In sum, mental scene construction according to the semantic account is lacking an explicit time and incentive model, both of which are integral parts of the MDP interpretation of DMN function.

### 4.3 Sentinel hypothesis

Regions of the DMN have been proposed to process the experienced or expected relevance of environment cues (Montague et al., 2006). Processing self-relevant information was perhaps the first functional account that was proposed for the DMN (Gusnard et al., 2001; Raichle et al., 2001). Since then, many investigators have speculated that neural activity in the DMN may reflect the brain’s continuous tracking of relevance in the environment, such as spotting predators, as an advantageous evolutionary adaptation (Buckner et al., 2008; Hahn et al., 2007). According to this cognitive account, the human brain’s baseline maintains a “radar” function to detect subjectively relevant cues and unexpected events in the environment. Propositions of a sentinel function to underlie DMN activity have however seldom detailed the mechanisms of how attention and memory resources are exactly reallocated when encountering a self-relevant environmental stimulus. Instead, in the present MDP account, promising action trajectories are recursively explored by the human DMN. Conversely, certain branches of candidate action trajectories are detected to be less worthy to get explored. This mechanism, expressed by the Bellman equation, directly implies stratified allocation of attention and working memory load over relevant cues and events in the environment. Further, our account provides a parsimonious explanation for the consistently observed DMN implication in certain goal-directed experimental tasks and in task-unconstrained mind-wandering (Smith et al., 2009; Bzdok et al., 2016b). Both environment-detached and environment-engaged cognitive processes may entail DMN recruitment if real or imagined experience is processed, manipulated, and used in service of organism control. During active engagement in tasks, the policy and value estimates may be updated to optimize especially short-term action. At passive rest, these parameter updates may improve especially mid and long-term action. This horizon of the agent is expressed in the *γ* parameter in the MDP account. We thus provide answers for the currently unsettled question why the involvement of the same neurobiological brain circuit (i.e., DMN) has been documented for specific task performances and baseline house-keeping’ functions.

In particular, environmental cues that are especially important for humans are frequently of social nature. This may not be surprising given that the complexity of the social systems is likely to be a human-defining property (Tomasello, 2009). According to the “social brain hypothesis”, the human brain has especially been shaped for forming and maintaining increasingly complex social systems, which allows solving ecological problems by means of social relationships (Whiten and Byrne, 1988). In fact, social topics probably amount to roughly two thirds of human everyday communication (Dunbar et al., 1997). Mind-wandering at daytime and dreams during sleep are also rich in stories about people and the complex interactions between them. In line with this, DMN activity was advocated to be specialized in continuous processing of social information as a physiological baseline of human brain function (Schilbach et al., 2008). This view was later challenged by observing analogues of the DMN in monkeys (Mantini et al., 2011), cats (Popa et al., 2009), and rats (Lu et al., 2012), three species with social capacities that are supposedly less advanced than in humans.

Further, the principal connectivity gradient in the cortex appears to be greatly expanded in humans compared to monkeys, suggesting a phylogenetically conserved axis of cortical expansion with the DMN emerging at the extreme end in humans (Margulies et al., 2016a). Computational models of dyadic whole-brain dynamics demonstrated how the human connectivity topology, on top of facilitating processing at the intra-individual level, can explain our propensity to coordinate through sensorimotor loops with others at the inter-individual level (Dumas et al., 2012). The DMN is moreover largely overlapping with neural networks associated with higher-level social processes (Schilbach et al., 2012). For instance, the vmPFC, PMC, and RTPJ together may play a key role in bridging the gap between self and other by integrating low-level embodied processes within higher level inference-based mentalizing (Lombardo et al., 2009).

Rather than functional specificity for processing social information, the present MDP account can parsimoniously incorporate the dominance of social content in human mental activity as high value function estimates for information about humans (Baker et al., 2009; Kampe et al., 2001; Krienen et al., 2010). The DMN may thus modulate reward processing in the human agent in a way that prioritizes appraisal of and action towards social contexts, without excluding relevance of environmental cues of the physical world. In sum, our account on the DMN directly implies its previously proposed “sentinel” function of monitoring the environment for self-relevant information in general and inherently accommodates the importance of social environmental cues as a special case.

### 4.4 The free-energy principle and active inference

According to theories of the *free-energy principle* (FEP) and *active inference* (Friston, 2010; Friston et al., 2009; Dayan et al., 1995), the brain corresponds to a biomechanical reasoning engine. It is dedicated to minimizing the long-term average of surprise: the log-likelihood of the observed sensory input more precisely, an upper bound thereof relative to the expectations about the external world derived from internal representations. The brain would continuously generate hypothetical explanations of the world and predict its sensory input x (analogous to the state-action (*s,a*) pair in an MDP framework). However, surprise is challenging to optimize numerically because we need to sum over all hidden causes z of the sensations (an intractable problem). Instead, FEP therefore minimizes an upper-bound on surprise given by

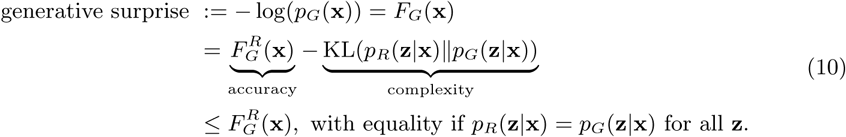

where

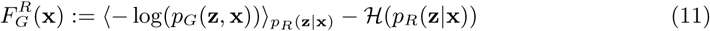

is the *free energy.* Here, the angular brackets denote the *expectation* of the joint negative log-likelihood — log(*p*_*G*_(z,x)) w.r.t the recognition density *p*_*R*_(z |x), *H* is the *entropy* functional defined by *H*(*p*) ≔ — Σ_z_ *p*(z)log(*p*(z),while KL(.‖.) is the usual *Kullback-Leibler (KL) divergence* (also known as *relative entropy*) defined by KL(*p*‖*q*) ≔ Σ_z_*p*(z)log(*p*(z)/*q*(z)) ≥ 0, which is a measure of how different two probability distributions are. In this framework, the goal of the agent is then to iteratively refine the generative model *p* _G_ and the recognition model *p*_*R*_ so as to minimize the free energy 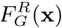over sensory input x.

Importantly,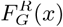gets low in the following cases:

- *p*_*R*_(z|x) puts a lot of mass on configurations (z,x) which are *pG* likely, **and**
- *p*_*R*_(z|x) is as uniform as possible (i.e have high entropy), so as not to concentrate all its mass on a small subset of possible causes for the sensation x.

Despite its popularity, criticism against the FEP has been voiced over the years, some of which is outlined in the following. The main algorithm for minimizing free energy 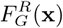 is the *wake-sleep algorithm* (Dayan et al., 1995). As these authors noted, a crucial drawback of the wake-sleep algorithm (and therefore of theories like the FEP (Friston, 2010)) is that it involves a pair of forward (generation) and backward (recognition) models *p*_*G*_ and *p*_*R*_ that together does not correspond to optimization of (a bound of) the marginal likelihood because KL divergence is not symmetric in its arguments.

These considerations render the brain less likely to implement the wake-sleep algorithm or a variant thereof. More recently, *variational auto-encoders* (VAEs) (Kingma and Welling, 2013) emerged that may provide an efficient alternative to the wake-sleep algorithm. VAEs overcome a number of the technical limits of the wake-sleep algorithm by using a reparametrization maneuver, which makes it possible to do differential calculus on random sampling procedures without exploding variance. As a result, unlike the wake-sleep algorithm for minimizing free energy, VAEs can be efficiently trained via back-propagation of prediction errors.

The difference between the FEP and the MDP account may be further clarified by a thought experiment. Since theories based on the FEP (Friston, 2010; Friston et al., 2009) conceptualize ongoing behavior in an organism to be geared towards the surprise-minimizing goal, an organism entering a dark room (Fig. 6) would remain trapped in this location because its sensory inputs are perfectly predictable given the environmental state (Friston et al., 2012). However, such a behavior is seldom observed in humans in the real world. In a dark room, the intelligent agents would search for light sources by explore the surroundings or aim to exit the room. Defenders of the FEP have retorted by advancing the “full package” (Friston et al., 2012): FEP is proposed to be multi-scale and there would be a meta-scale at which the organism would be surprised by such a lack of surprise. According to this argument, a dark room would paradoxically correspond to a state of particularly high relevance. Driven by the surprise-minimization objective, the FEP agent would eventually bootstrap itself out of such saddle points to explore more interesting parts of the environment. In contrast, an organism operating under our RL-based theory would inevitably identify the sensory-stimulus-deprived room as a local minimum. Indeed, hippocampal experience replay (see 3.2.3) could serve to sample memories or fantasies of alternative situations with reward structure. Such artificially generated internal sensory input subserved by the DMN can entice the organism to explore the room, for instance by looking for and using the light switch or finding the room exit.

**Fig. 6.**
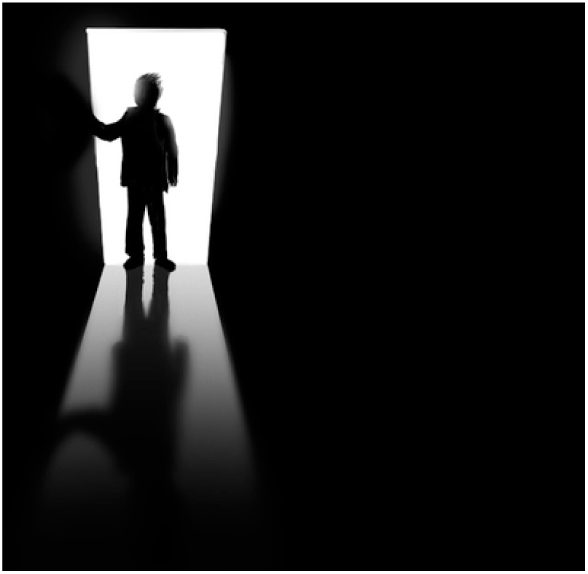
The dark room experiment. An intelligent agent situated in a light-deprived closed space is used as a thought experiment for the complete absence of external sensory input.

We finally note that FEP and active inference can be reframed in terms of our model-free RL framework. This becomes possible by recasting the Q-value function (i.e., expected long-term reward) maximized by the DMN to correspond to negative surprise, that is, the log-likehood of current sensory priors the agent has about the world. More explicitly, this corresponds to using free-energy as a *Q*-value approximator for the MDP such that

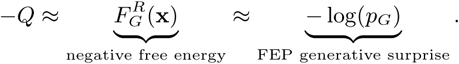

Such a surprise-guided RL scheme has previously been advocated under the equivalent framework of energy-based RL (Sallans and Hinton, 2004; Elfwing et al., 2016) and information compression (Schmidhuber, 2010; Mohamed and Rezende, 2015). Nevertheless, minimization of surprise quantities alone may be insufficient to explain the diversity of behaviors that humans and other intelligent animals can perform.

## 5 Conclusion

Which brain function could be important enough for the existence and survival of the human species to justify constantly high energy costs? MDPs motivate an attractive formal account of how the human association cortex can be thought to implement multi-sensory representation and high-level decision-making to optimize the organism’s intervention on the world. This idealized process model accommodates a number of previous observations from neuroscience studies on the DMN by simple but non-trivial mechanisms. Viewed as a Markovian sequential decision process, human behavior unfolds by inferring cumulative reward outcomes from hypothetical action cascades and extrapolation from past experience to upcoming events for guiding behavior in the present. MDPs also provide a formalism how opportunity in the environment can be deconstructed, evaluated, and exploited when confronted with challenging interdependent decisions. This functional interpretation may well be compatible with the DMN’s poorly understood involvement across autobiographical memory recall, problem solving, abstract reasoning, social cognition, as well as delay discounting and self prospection into the future. Improvement of the internal world representation by injecting stochasticity into the recall of past actions and inference of action outcomes may explain why highly accurate memories have been disfavored in human evolution and why human creativity may be adaptive.

A major hurdle in understanding DMN activity from cognitive brain-imaging studies has been its similar neural engagement in different time-scales: thinking about the past (e.g., autobiographical memory), thinking about hypothetical presents (e.g., daytime mind-wandering), and thinking about anticipated scenarios (e.g., delay discounting). The MDP account of DMN function offers a natural integration of a-priori diverging neural processes into a common framework. It is an important advantage of the proposed artificial intelligence perspective on DMN biology that it is practically computable and readily motivates neuroscientific hypotheses that can be put to the test in future research. Neuroscience experiments on the DMN should be designed that operationalize the set of action, value, and state variables governing the behavior of intelligent RL agents. At the least, we propose an alternative vocabulary to describe, contextualize, and interpret experimental findings in neuroscience studies on higher-level cognition. Ultimately, neural processes in the DMN may realize a holistic integration ranging from real experience over purposeful dreams to predicted futures to continuously refine the organism’s fate.

